# Sex chromosome dosage compensation in *Heliconius* butterflies: global yet still incomplete?

**DOI:** 10.1101/016675

**Authors:** James R. Walters, Thomas J. Hardcastle, Chris D. Jiggins

## Abstract

The evolution of heterogametic sex chromosome is often – but not always – accompanied by the evolution of dosage compensating mechanisms that mitigate the impact of sex-specific gene dosage on levels of gene expression. One emerging view of this process is that such mechanisms may only evolve in male-heterogametic (XY) species but not in female-heterogametic (ZW) species, which will consequently exhibit “incomplete” sex chromosome dosage compensation. However, recent results suggest that at least some Lepidoptera (moths and butterflies) may prove to be an exception to this prediction. Studies in bombycoid moths indicate the presence of a chromosome-wide epigenetic mechanism that effectively balances Z chromosome gene expression between the sexes by reducing Z-linked expression in males. In contrast, strong sex chromosome dosage effects without any reduction in male Z-linked expression were previously reported in a pyralid moth, suggesting a lack of any such dosage compensating mechanism. Here we report an analysis of sex chromosome dosage compensation in *Heliconius* butterflies, sampling multiple individuals for several different adult tissues (head, abdomen, leg, mouth, and antennae). Methodologically, we introduce a novel application of linear mixed-effects models to assess dosage compensation, offering a unified statistical framework that can estimate effects specific to chromosome, to sex, and their interactions (*i.e.,* a dosage effect). Our results show substantially reduced Z-linked expression relative to autosomes in both sexes, as previously observed in bombycoid moths. This observation is consistent with an increasing body of evidence that some lepidopteran species possess an epigenetic dosage compensating mechanism that reduces Z chromosome expression in males to levels comparable with females. However, this mechanism appears to be imperfect in *Heliconius*, resulting in a modest dosage effect that produces an average 5-20% increase in male expression relative to females on the Z chromosome, depending on the tissue. Thus our results in *Heliconius* reflect a mixture of previous patterns reported for Lepidoptera. In *Heliconius,* a moderate pattern of “incomplete” dosage compensation persists apparently despite the presence of an epigenetic dosage compensating mechanism. The chromosomal distributions of sex-biased genes show an excess of male-biased and a dearth of female-biased genes on the Z chromosome relative to autosomes, consistent with predictions of sexually antagonistic evolution.

## Introduction

Dosage compensation is a gene-regulatory mechanism that equalizes levels of gene expression in response to differences in gene dose (i.e. copy number). Without dosage compensation, changes in gene dose can substantially affect gene expression, potentially resulting in detrimental effects on finely-tuned gene networks (Birchler et al. 2001). For this reason, it was long assumed that the evolution of a chromosome-wide dosage compensating mechanism was an essential step in the evolution of heteromorphic sex chromosomes (Vicoso & Bachtrog 2009; Disteche 2012; Mank 2013).

Heteromorphic sex chromosomes typically evolve from homologous autosomes that acquire a sex-determining locus, accumulate sexually-antagonistic alleles, and suppress recombination (B Charlesworth 1996; B Charlesworth & D Charlesworth 2000; Bachtrog 2006). Eventually, substantial gene loss and degeneration occurs on the nascent Y chromosome (or W in female-heterogametic ZW taxa) (Rice 1984; B Charlesworth & D Charlesworth 2000; Bachtrog 2013). The erosion of genes from the Y/W presents the problem of balancing gene expression with the autosomes. Monosomy of the X/Z in one sex means the dose (*i.e.,* copy number) of most sex-linked genes differs by half between the sexes. If the resulting sex-linked gene expression becomes similarly unbalanced, degradation of the Y/W could impose a substantial fitness cost for the heterogametic sex due to impaired dosage-sensitive interactions with autosomal loci (Ohno 1967; Mank 2009; Pessia et al. 2013). It is thus predicted that a global dosage-compensating mechanism should evolve to balance the expression of X/Z loci relative to autosomes as the Y/W degrades (Ohno 1967; Brian Charlesworth 1978; Mank 2013; Veitia et al. 2015). This process should also result, indirectly, in balanced gene expression between the sexes for the X/Z. One important working assumption in this framework is that average expression is approximately equal across autosomes, therefore *complete* dosage compensation should yield X:A (or Z:A) expression ratios of ∼1 in both sexes (Mank 2009; 2013; Nguyen & Disteche 2006; Vicoso & Bachtrog 2011b; Smith et al. 2014; Walters & Hardcastle 2011).

Efforts to evaluate this hypothesis have expanded greatly in the last decade with the application of genome-wide expression analyses via microarray or RNA-seq, and have yielded several unexpected results. The history and contemporary findings of research on sex chromosome dosage compensation are extensively reviewed in several recent publications (Disteche 2012; Mank 2013; Pessia et al. 2013; Ferrari et al. 2014). Here we briefly highlight details particularly relevant to our current results. Where investigated during the pre-genomic decades, the limited results obtained tended to support theoretical predictions. Initial genome-wide investigations using microarrays in established model organisms such as mouse, humans, fruit flies, and nematodes – all male heterogametic species – yielded patterns consistent with global sex chromosome dosage compensation. As predicted, average expression on the X was comparable to autosomes (X:A ∼ 1) in both sexes and the canonical view of dosage compensation and sex chromosome evolution appeared robust (Hamada 2005; Gupta et al. 2006; Nguyen & Disteche 2006). Moreover, the recognition that distinct molecular mechanism underlay dosage compensation in flies, worms, and humans added further support to the universality of dosage compensation evolving concomitantly with differentiated sex chromosomes (Deng et al. 2011; Straub & Becker 2011). However, more recent research has added substantial complexity and controversy to the issue. In particular, evaluating dosage compensation in the context of ancestral expression levels indicates that eutherian mammals should be considered to have X:A expression ratio of ∼0.5 in both sexes (Julien et al. 2012; F Lin et al. 2012). Importantly, in all of these cases, sex-linked expression appears to be balanced between males and females (male:female ∼ 1 on the X) and there is no evidence of a gene dosage-effect on X chromosome expression.

Other striking exceptions to the canonical theory of dosage compensation were observed when investigations of sex chromosome dosage compensation expanded into novel taxa. Notably, several female-heterogametic taxa exhibit incomplete dosage compensation: Male birds, snakes, and schistosomes are homogametic (ZZ) with Z:A ∼ 1, but in females the Z:A ratio is significantly less than 1 (Itoh et al. 2007; Vicoso & Bachtrog 2011a; Vicoso, Emerson, et al. 2013a; Mank & Ellegren 2009; Naurin et al. 2011). The apparently dichotomous pattern of completely dosage compensated XY taxa versus incompletely compensated ZW species catalyzed strong suggestions and some nascent theory claiming that global, complete sex chromosome dosage compensation might be universally absent from ZW taxa and occur only XY species (Bachtrog et al. 2011; Mank 2013; Naurin et al. 2010; Vicoso, Emerson, et al. 2013a). We call this the *heterogametic dichotomy hypothesis*.

Despite the emerging evidence supporting the heterogametic dichotomy hypothesis, there is at least one conspicuous female-heterogametic taxon that may prove to be an exception: Lepidoptera (moths and butterflies). The status of sex chromosome dosage compensation in Lepidoptera is currently ambiguous, primarily due to inconsistent results reported by the few studies currently available. The first genome-wide assessment of sex chromosome dosage compensation in Lepidoptera was based on a microarray dataset in the silkmoth, *Bombyx mori* (Xia et al. 2007). An initial analysis of these data reported incomplete dosage compensation similar to other ZW taxa (Zha et al. 2009). However, analytical flaws were later identified in this initial effort and a subsequent reanalysis indicated the Z:A ratio was equal in males and females, offering evidence for global, complete sex chromosome dosage compensation (Walters & Hardcastle 2011). Intriguingly, this reanalysis further revealed a Z:A ratio of ∼0.7, significantly less than one. This result is not anticipated by current theory concerning sex chromosome evolution. Additional evidence for complete sex chromosome dosage compensation in Lepidoptera was more recently reported in another bombycoid moth, *Manduca sexta* (tobacco hormworm), using RNA-seq (Smith et al. 2014). Again the Z:A ratio was equal between sexes, but in this case the Z:A ratio was only marginally less then 1, with significance depending on filtering thresholds. In stark contrast to results from Bombycoid moths, RNA-seq analysis of the Pyralid moth *Plodia interpunctella* (Indian meal moth) showed no evidence for dosage compensation, with female Z:A∼0.5 while male Z:A was ∼ 1 (Harrison et al. 2012). This is the largest magnitude of sex chromosome dosage effect yet reported, exceeding patterns observed in birds and snakes (Itoh et al. 2007; Vicoso & Bachtrog 2011a; Vicoso, Emerson, et al. 2013a; Mank & Ellegren 2009; Naurin et al. 2011).

These studies differed considerably in what tissues or body parts were assayed. The *B. mori* microarray data included samples from 10 different larval body parts (Xia et al. 2007; Walters & Hardcastle 2011). The *M. sexta* study sampled only adult heads while pools of whole adult *P. interpunctella* were used (Harrison et al. 2012; Smith et al. 2014). These latter two studies both constructed *de novo* transcriptome assemblies and assigned chromosomal linkage based on contig homology to *B. mori.* This approach of assigning Z-linkage is seemingly robust because synteny is highly conserved in Lepidoptera (Pringle et al. 2007; Heliconius Genome Consortium 2012; Yue et al. 2013; Ahola et al. 2014).

However, only the *B. mori* dataset, which included isolated gonads, provided any opportunity to assess patterns of dosage compensation separately in somatic and reproductive tissues. It is well-established that, at least in *B. mori*, the Z chromosome is enriched for highly-expressed, testes-specific genes but depleted of ovary-specific transcripts (Arunkumar et al. 2009; Suetsugu et al. 2013). Thus, inclusion of gonadal tissue may substantially skew patterns of Z:A ratios to appear “uncompensated” even when somatic Z:A ratios are otherwise comparable between the sexes (Walters & Hardcastle 2011).

In this manuscript we report the first genomic analysis of sex chromosome dosage compensation in a butterfly. Using RNA-seq, we assay male and female gene expression in several body parts of *Heliconius melpomene* and its closely related sibling species, *H. cydno* (Martin et al. 2013; Quek et al. 2010). The existence of a complete reference genome and linkage map for *H. melpomene* facilitates a nuanced inference of chromosome and sex specific effects on gene expression, which we achieve through the novel application of mixed-effects linear models to analyze dosage compensation (Heliconius Genome Consortium 2012). Our results show substantially reduced Z-linked expression relative to autosomes in both sexes, but also modest dosage effect on the Z chromosome, and thus reflect a mixture of previous patterns reported for Lepidoptera.

## Methods

### Samples and Sequencing

Two groups of RNA-seq data were used to estimate gene expression. First, we used the paired-end data sequenced from *H. melpomene* generated by Briscoe et al. (Briscoe et al. 2013) ArrayExpress ID: E-TAB-1500. This data set includes three male and three female samples from mouth, leg, and antennae. Additionally, we generated new RNA-seq data from adult head (excluding antennae) and abdomen. For these samples, *Heliconius* butterflies were reared in large insectaries in Gamboa, Panama. Insectary populations were recently established from local natural populations. Males and females were kept separate after eclosion and aged 6 days before collection to allow reproductive tissues to develop (Dunlap-Pianka et al. 1977); all samples were virgins. Head and abdomen tissues were collected into RNAlater and stored frozen before RNA purification. Total RNA was extracted with TRIzol reagent (Invitrogen, Carlsbad, CA), purified using RNeasy columns (Qiagen, Valencia, CA), and treated with TURBO DNase (Life Technologies, Grand Island, NY) following manufacturer’s instructions. Messenger RNA was isolated from total RNA via poly-A pulldown and subsequently transformed into a cDNA library using the Illumina TruSeq sample preparation kits. Paired-end 100 basepair sequencing was performed on an Illumina HiSeq.

### Read mapping and normalization

Read mapping and estimation of fragment counts per gene were performed using RSEM (v1.2.11) running Bowtie2 (v2.1.0) (Li & Dewey 2011; Langmead & Salzberg 2012). Reads were mapped as paired-end data, with the first 9 bp and the final 40 bp trimmed before mapping to remove low and variable-quality bases. All subsequent statistical analyses were performed using R and BioConductor, especially the baySeq package for assessing differential expression (Hardcastle & Kelly 2010; R Development Core Team 2014). The library scaling factors were calculated as the sum of non-zero gene expression levels below the 75th percentile of gene expression, following the example of Hardcastle et al. (2012) as an amendment to Bullard *et al.* (2010). As there is potential for substantial Z chromosome effects on gene expression, only known autosomal loci were considered when calculating library scaling factors. For non-parametric statistical analyses, we further normalized expression levels as fragments per kilobase per million mapped reads (FPKM).

Failing to remove transcriptionally inactive genes from RNA-seq data sets assayed for sex chromosome dosage compensation can result in problematic biases, yet the most appropriate filtering method to apply is not well-established (Xiong et al. 2010; Kharchenko et al. 2011; Jue et al. 2013; Smith et al. 2014). Here we employ a probabilistic approach to assessing whether a given locus is expressed. The baySeq framework calculates the posterior likelihood (ranging from 0 to 1) that a given locus has no true expression. (i.e., any observed reads should be considered “noise”, not signal) (Hardcastle 2014). We primarily report results filtered at a value of 0.5, but results are comparable across a range thresholds tested, from 0.25 to 0.9 (see Supplementary Information)

### Assessing Z chromosome and dosage effects: non-parametric statistics

To test for Z chromosome dosage effects, we compared the expression of Z-linked and autosomal loci in males and females. Greater than 80% of predicted coding loci have been mapped to the 21 chromosomes of *H. melpomene*, with Z-linkage validated or corrected based on sex-specific genome sequencing coverage as reported in Martin *et al.* 2013 (Martin et al. 2013; Heliconius Genome Consortium 2012). For each body part sampled, replicates were averaged by sex to give mean male and female FPKM values for each locus. Within each sex, Z-linked versus autosomal (Z:A) expression was compared, with differences in average expression evaluated via Mann-Whitey U test (MWU). The average male:female (M:F) expression ratio of Z-linked versus autosomal loci was also compared via MWU. Only loci actively expressed in both sexes were included when analyzing M:F ratios.

We further explored the effects of Z chromosome dosage by comparing the average expression of Z-linked genes in males and females, split by quartiles of expression magnitude. Within quartile, differences between sex in average Z-linked expression was tested via MWU. Only loci actively expressed in both sexes were included in this analysis. This analysis closely resembles an analysis performed by Harrison *et al.* (2012) aimed at assessing how Z chromosome dosage effects depend on expression magnitude. However, we have slightly modified the analysis to avoid a bias we believe is inherent in the analysis as originally performed by Harrison *et al.* (2012). Rather than basing expression quartiles solely on male expression as was previously done, we calculated quartiles based on the maximum of male or female expression for each locus. The reasoning for this modification and the potential biases arising from ranking genes using data from only one sex is provided in the Appendix.

### Assessing sex-chromosome and dosage effects: Linear modeling of expression levels

In addition to the application of non-parametric MWU tests, as is typically employed for investigations of sex-chromosome dosage compensation, we implemented a linear modeling framework to test for dosage and Z-specific effects on gene expression.

The count data (after filtering) were fitted via maximum likelihood methods (Bates et al. 2014) to a generalized linear mixed-effects model assuming a Poisson distribution and a log linkage function, using library scaling factor and gene length as offsets. We fit a series of models using various effects of chromosome, Z-linkage and sex, together with relevant interactions between these effects. In all models, a per gene random effect was applied over sex, simultaneously allowing for variation in individual gene expression and differential expression of genes between the sexes.

Using this modeling framework, we first tested for a global effect of Z-linkage on expression by comparing a model with both **sex** and **Zlink** as fixed effects (but no interaction)

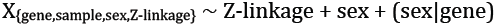

versus a model with **sex** as the sole fixed effect.

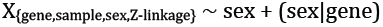

We then additionally tested for sex-specific effect on Z-expression (*i.e.,* a dosage effect), by fitting the full model

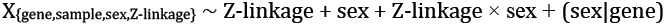

to one without the interaction term.

### Chromosomal distribution of sex-biased genes

Genes differentially expressed between males and females were identified using baySeq (Hardcastle & Kelly 2010). We applied a false-discovery rate of 0.05 and required at least a 1.5-fold change in expression between sexes. To examine the chromosomal distribution of sex-biased genes, we counted the number of male, female, and unbiased genes among all actively expressed genes on each chromosome and also the unmapped scaffolds not yet assigned to chromosome. Gene activity was based on the probabilistic criteria, and assessed independently in males and females, so a gene expressed in males but not females was counted as male biased (and vice versa). Differences between the Z and autosomes in proportion of sex-biased genes were tested using a Fisher’s exact test.

### Results

#### Sequencing and read mapping

Data sets newly generated for this project were considerably larger that those generated by Briscoe et al. (2013). For the 40 samples sequenced here, the mean number of total reads sequenced was ∼130 M, ranging from ∼30 M to ∼ 284 M reads. The 18 samples from Briscoe et al. (2013) gave a mean of ∼23 M, ranging from ∼8 M to ∼46 M reads. The proportion of reads aligned with RSEM was also generally higher for the abdomen and head samples compared to the Briscoe et al samples from antennae, mouth, and leg. The 40 abdomen and head samples yielded a mean of 49% aligned, ranging from 21 to 65%. Mean percent mapping in the 18 Briscoe et al. (2013) samples was 37%, ranging from 9 to 55%. A complete summary of read counts and alignment statistics calculated with Picard’s CollectAlignmentSummaryMetrics utility (http://picard.sourcefource.net) is provided in supplementary Table S1. Newly generated data sets for *H. melpomene* and *H. cydno* head and abdomen are available from the Sequencing Read Archive (SRA) project: XXXXX.

### Z-chromosome to autosome comparisons

For both sexes, all body parts in both species yielded average Z expression significantly lower than autosomal expression, as is evident from both linear modeling and non-parametric analyses (Fig 1 and Tables 1 & 2). Comparing median expression, the Z:A ratio ranged from about 0.5 to 0.75; ratios based on mean values are even lower. Linear modeling of these data showed Z-linkage had a significant negative effect on gene expression relative to autosomal average (Table 2). Average expression of individual autosomes varied somewhat in both sexes, but in nearly all cases the median Z expression was lower than any autosome (supplementary Fig 1)

**Fig 1.**
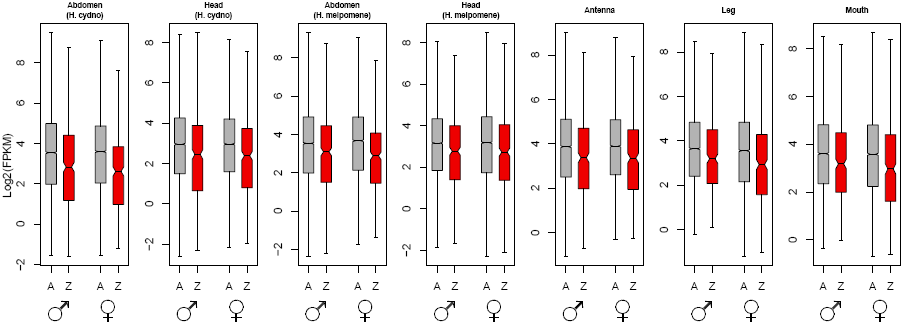
Distributions of log2-transformed gene expression levels (FPKM) for the Z-chromosome (Z; red) and autosomes (A; grey) in male and female *Heliconius* butterflies in several tissues. Boxes indicate the interquartile range (IQR) around the median (black bar) and whiskers extend to 1.5 times the IQR; outlier points beyond 1.5 IQR are not shown. Non-parametric tests as well as linear modeling indicate a significant reduction in Z-chromosome expression relative to autosomes in both sexes for all tissues (see Tables 1 & 2). Data were filtered with a null expression likelihood of 0.5 and are representative of results using a range of likelihood thresholds (supplementary Table 2).

**Table 1.** Summary of average expression of Z-linked and autosomal loci across several tissues in *Heliconius* butterflies. Results reflect data ing with a null expression likelihood of 0.5 and are representative of results using a range of likelihood thresholds (supplementary Table S2)

| Statistic | Abdomen<br>(H. cydno) |  | Head<br>(H. cydno) |  | Abdomen<br>(H. melpomene) |  | Head<br>(H. melpomene) |  | Antenna |  | Leg |  | Mouth |  |
| --- | --- | --- | --- | --- | --- | --- | --- | --- | --- | --- | --- | --- | --- | --- |
|  | Male | Female | Male | Female | Male | Female | Male | Female | Male | Female | Male | Female | Male | Female |
| Mean Z-linked FPKM | 40.66 | 26.15 | 29.27 | 23.78 | 36.63 | 29.6 | 28.14 | 29.18 | 30.32 | 27.3 | 47.99 | 41.65 | 40.74 | 35.07 |
| Mean autosomal FPKM | 85.45 | 54.81 | 70.64 | 53.6 | 63.42 | 53.33 | 54.2 | 68.43 | 75.66 | 67.84 | 63.3 | 59.02 | 71.77 | 68.06 |
| Z:A ratio of mean expression | 0.4758 | 0.4771 | 0.4144 | 0.4437 | 0.5776 | 0.555 | 0.5192 | 0.4264 | 0.4007 | 0.4024 | 0.7581 | 0.7057 | 0.5676 | 0.5153 |
| Median Z-linked FPKM | 7.15 | 6.15 | 5.48 | 5.23 | 8.53 | 7.38 | 6.8 | 6.59 | 10.38 | 10.26 | 9.12 | 7.59 | 9.03 | 7.81 |
| Median autosomal FPKM | 11.67 | 12.12 | 7.78 | 7.82 | 11.6 | 12.71 | 8.93 | 9.03 | 14.65 | 14.93 | 12.43 | 11.69 | 12.13 | 11.96 |
| Z:A ratio of median expression | 0.6127 | 0.5074 | 0.7044 | 0.6688 | 0.7353 | 0.5806 | 0.7615 | 0.7298 | 0.7085 | 0.6872 | 0.7337 | 0.6493 | 0.7444 | 0.653 |
| MWU p-value:<br>Autosomal ≠ Z-linked | <.0001 | <.0001 | <.0001 | <.0001 | <.0001 | <.0001 | .0007 | .0001 | <.0001 | <.0001 | .0003 | <.0001 | .0002 | <.0001 |
| No. expressed Z-linked loci | 481 | 403 | 443 | 429 | 506 | 417 | 449 | 447 | 392 | 394 | 357 | 385 | 361 | 390 |
| No. expressed autosomal loci | 10638 | 9758 | 10272 | 10050 | 11020 | 10014 | 10004 | 10383 | 9296 | 9318 | 8548 | 9239 | 8792 | 9243 |

Notably, in several body parts sampled, the Z:A ratio was slightly greater in males than in females (Table 1). This apparent sex-effect on Z-chromosome expression was statistically confirmed with linear modeling. All tissues except antennae showed a significant interaction between sex and Z-linked expression, with greater male Z-linked expression (Table 2). This interaction can be interpreted as dosage effect of the Z chromosome on average gene expression. Note, however, that the magnitude of this interaction (dosage) effect is distinctly less than the effect of the Z-chromosome on gene expression levels.

### Male:Female expression ratios

Consistent with the dosage effect observed in the linear models, the Z chromosome showed a modest but consistent male bias in expression relative to autosomes in all samples (Figure 2 and Table 3). The distribution of M:F expression ratios was significantly greater on the Z (MWU p < 0.05 after bonferroni correction) in all samples except antennae. The average magnitude of this Z-linked male expression bias ranged from 5-20% relative to autosomes based on the median M:F expression ratios (Table 2).

**Fig 2.**
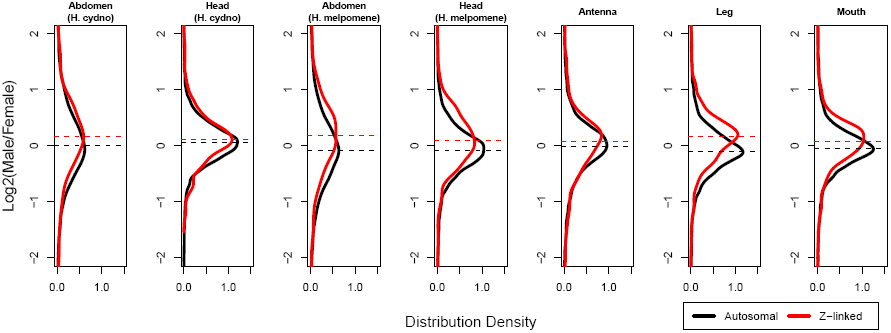
Distribution density plots of log2(Male:Female) expression ratios for the Z-chromosome (Z; red) and autosomes (A; black) in male and female *Heliconius* butterflies in several tissues. Dashed lines indicate median values. Non-parametric tests and linear modeling indicate all tissues but antenna show a modest but significant sex-chromosome dosage effect, visualized here as a shift towards male-biased expression among Z-linked loci (see Tables 2 & 3). Antenna also shows this pattern but the effect is not statistically significant.

**Table 2.** Linear modeling analysis of Z-chromosome and dosage effects on gene expression levels across several tissues in *Heliconius* butterflies. Results reflect data filtering with a null expression likelihood of 0.5 and are representative of results using a range of likelihood sholds (supplementary Table S3)

|  | Z-linkage only |  |  | Z-linkage x sex interaction (Dosage effect) |  |  |
| --- | --- | --- | --- | --- | --- | --- |
| Sample | Z-linkage Effect Size | ANOVA p-value |  | Z-linkage Effect Size | Interaction (Dosage) Effect Size | ANOVA p-value |
| Abdomen (H. cydno) | -0.56 | 2.16E-14 |  | -0.65878 | 0.212 | 2.57E-05 |
| Head (H. cydno) | -0.431 | 3.02E-08 |  | -0.40678 | 0.0642 | 7.66E-08 |
| Abdomen (H. melpomene) | -0.385 | 1.53E-07 |  | -0.47591 | 0.189 | 0.0001 |
| Head (H. melpomene) | -0.263 | 0.00030 |  | -0.4581 | 0.289 | 6.90E-27 |
| Antenna | -0.371 | 2.35E-07 |  | -0.37173 | 0.0307 | 0.224 |
| Leg | -0.275 | 0.00047 |  | -0.36309 | 0.169 | 1.47E-14 |
| Mouth | -0.340 | 6.72E-06 |  | -0.36206 | 0.0615 | 0.00227 |

**Table 3.** Summary of average Male:Female gene expression ratios for Z-linked and autosomal loci in *Heliconius* butterflies for several es. Results reflect data filtering with a null expression likelihood of 0.5 and are representative of results using a range of likelihood sholds (supplementary Table S4).

| Tissue | No. Z-linked loci | No. Autosomal loci | Mean Z-linked log2(M:F) | Mean autosomal log2(M:F) | Median Z – linked log2(M:F) | Mean autosomal log2(M:F) | Z:A ratio of means (Not Log2) | Z:A ratio of medians (Not Log2) | MWU p-value: Autosomal ≠ Z-linked |
| --- | --- | --- | --- | --- | --- | --- | --- | --- | --- |
| Abdomen (H. cydno) | 387 | 9539 | 0.3553 | 0.1101 | 0.1683 | -0.0009 | 1.1853 | 1.1244 | <.0001 |
| Head (H. cydno) | 425 | 9972 | 0.1173 | 0.0584 | 0.116 | 0.0543 | 1.0417 | 1.0437 | 0.0034 |
| Abdomen (H. melpomene) | 408 | 9859 | 0.2418 | 0.0106 | 0.1904 | -0.0788 | 1.1738 | 1.2051 | <.0001 |
| Head (H. melpomene) | 433 | 9909 | 0.0847 | -0.1372 | 0.1002 | -0.0904 | 1.1662 | 1.1412 | <.0001 |
| Antenna | 378 | 8957 | 0.0115 | -0.039 | 0.0731 | -0.009 | 1.0356 | 1.0586 | 0.021 |
| Leg | 354 | 8457 | 0.1272 | -0.0912 | 0.1731 | -0.097 | 1.1634 | 1.2059 | <.0001 |
| Mouth | 356 | 8636 | 0.003 | -0.0741 | 0.0811 | -0.0561 | 1.0549 | 1.0997 | <.0001 |

### Quartile analysis of Z expression

Comparing Z-linked expression between sexes split by quartiles of expression showed no obvious pattern of discrepancy in male versus female Z-linked expression for tissues other than abdomen (Figure 3; supplemental Figure S2). In both abdomen samples, male expression was significantly greater than female in the fourth (highest expression) quartile, assuming a bonferroni correction for four tests.

**Fig 3.**
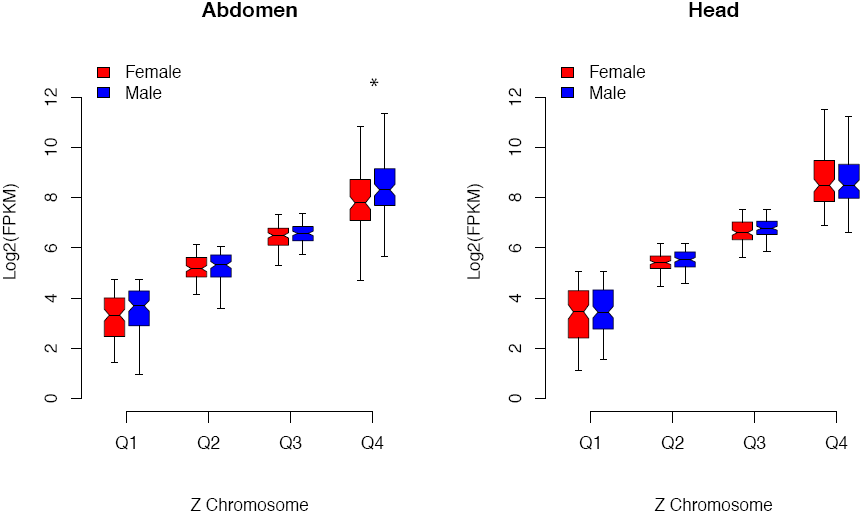
Comparison of male (blue) and female (red) expression levels for Z-linked loci, divided into quartiles based on the maximum of male or female expression for *H. melpomene* head and abdomen. Boxes indicate the interquartile range (IQR) around the median (black bar) and whiskers extend to 1.5 times the IQR. An asterisk (*) indicates significant difference in average expression (bonferroni corrected MWU p-value < 0.05). Plots for all seven tissue samples are given in supplemental Figure S2.

### Chromosomal distribution of sex-biased genes

The amount of sex-biased gene expression differed substantially between body parts. Tissues other than in the abdomen showed almost no differential expression between sexes using the criteria we applied (Table 4). In contrast, roughly 30% of active genes in abdomen were differentially expressed. We therefore analyzed the chromosomal distribution of sex-biased genes only in the abdomen. We observed a significant difference in the proportions of sex-biased genes on the Z versus the autosomes in both *H. melpomene* and *H. cydno* (Fisher’s exact test, p-value ≪ 0.001). Figure 4 shows that the Z chromosome is distinctly enriched for male-biased and has a paucity of female-biased genes.

**Fig 4.**
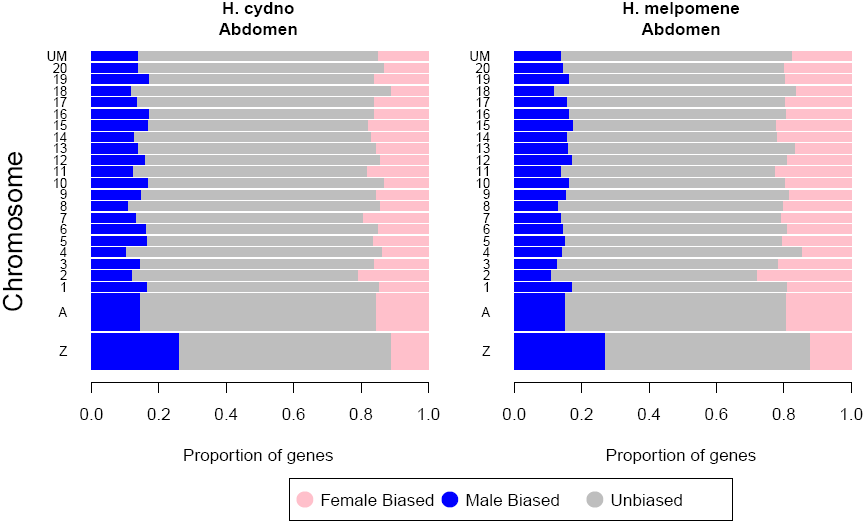
Proportions of sex-biased genes across the chromosomes in abdomens for *H. cydno* and *H. melpomene*. A gene was considered sex-biased if significantly differentially expressed between males and females with FDR < 0.05 and a minimum fold-change of 1.5. The two larger bars at the bottom represent the Z chromosome individually and the combined results across autosomes (A). Smaller bars (1-20) represent data for individual chromosomes and “UM” indicates genes not mapped to chromosome.

**Table 4.** Counts of sex-biased and unbiased genes (FDR < 0.05 and a minimum fold-change of 1.5) on the Z chromosome and autosomes (A) veral tissues of *Heliconius* butterflies.

| Expression<br>Category | Abdomen<br>(H. cydno) |  | Head<br>(H. cydno) |  | Abdomen<br>(H. melpomene) |  | Head<br>(H. melpomene) |  | Antenna |  | Leg |  | Mouth |  |
| --- | --- | --- | --- | --- | --- | --- | --- | --- | --- | --- | --- | --- | --- | --- |
|  | Z | A | Z | A | Z | A | Z | A | Z | A | Z | A | Z | A |
| Male bias | 114 | 1470 | 3 | 7 | 130 | 1590 | 0 | 0 | 0 | 0 | 0 | 0 | 0 | 0 |
| Unbiased | 275 | 7008 | 385 | 9389 | 290 | 6888 | 394 | 9354 | 319 | 8167 | 301 | 7717 | 295 | 7715 |
| Female bias | 48 | 1540 | 0 | 8 | 58 | 2025 | 0 | 3 | 0 | 3 | 0 | 0 | 0 | 0 |

## Discussion

### Dosage compensation

Patterns of sex chromosome dosage compensation in *Heliconius* butterflies reflect an interesting amalgam of previous results from Lepidoptera. Similarly to the bombycoid moths *B. mori* and *M. sexta*, *Heliconius* males show reduced expression of Z chromosome genes below autosomal expression levels (Walters & Hardcastle 2011; Smith et al. 2014). In the two bombycoid species, male Z expression was low enough to be comparable to the female Z; there was no detectable dosage effect on Z-linked expression. Yet despite having Z:A < 1 in both sexes, *Heliconius* shows a dosage effect such that the Z chromosome retains a modest but consistent male-bias in expression. This result in *Heliconius* echoes the very substantial dosage effects reported for *P. interpunctella*. Unfortunately, the magnitude of these *Heliconius* dosage effects cannot easily be contrasted with *P. interpunctella* because direct male:female expression ratios were not reported for that species (Harrison et al. 2012). However, the discrepancy in Z:A ratios between sexes of *P. interpunctella* is so large that we presume the dosage effects reflected in M:F expression ratios are far greater than we observe in *Heliconius*. Notably, no reduction in male Z-linked expression relative to autosomes was reported for *P. interpuctella*.

Thus in *Heliconius* we have further evidence that some Lepidoptera appear to have a molecular mechanism for globally reducing Z-linked expression in males that compensates for the difference in Z-chromosome dosage between sexes. Curiously, in *Heliconius* this mechanism appears to be operating imperfectly, resulting in a measurable dosage effect that could arguably be called “incomplete” dosage compensation. However, the 5-20% Z-linked male bias in *Heliconius* is distinctly less than the ∼50% bias typically reported for vertebrate ZW species (*e.g.,* snakes, birds) identified as having “incomplete” sex chromosome dosage compensation (Ellegren et al. 2007; Itoh et al. 2010; Vicoso, Emerson, et al. 2013a). In the case of the female-heterogametic vertebrates, it is assumed that dosage compensation is “incomplete” because no global epigenetic mechanism exists to offset dosage effects, in contrast to the epigenetic mechanisms known in fruit flies, nematodes, and mammals. In the case of *Heliconius*, we would argue that “incomplete” dosage compensation occurs despite a global mechanism operating to balance Z:A ratios between the sexes, in this case reducing male Z-linked expression similar to that in females.

Further evidence that a global mechanism mitigates dosage effects in *Heliconius* comes from the lack of detectable transcriptional saturation on the Z chromosome. None of the purely somatic tissues sampled showed an effect of expression magnitude on differences between male and female Z-linked expression. This result contrasts with *P. interpunctella*, where the male expression bias increased with expression level, suggesting substantial transcriptional saturation due to uncompensated Z chromosome dosage. However, in abdomens we did observe significantly greater expression in males for the highest expression quartile, reminiscent of the *P. interpunctella* result. While this could be regarded as the effect of transcriptional saturation due to gene imbalance (*i.e.,* a dosage effect), we tend to think it primarily reflects the unusually high expression levels of male-biased reproductive genes in the testes. If male-biased genes tend to have very high expression, this could produce the pattern observed. Robustly testing this hypothesis would require sequencing transcriptomes of gonads separately from somatic tissue; such data are currently not available in *Heliconius*. However, we note that in male abdomens, autosomal male-biased genes are expressed at significantly greater levels on average than unbiased or female biased genes (Kruskal-Wallis test, p ≪ 0.001; Supplemental Figure S3). In female abdomen, average expression of autosomal sex-biased genes does not differ from unbiased genes (Kruskal-Wallis test, N.S.). These patterns are consistent with our hypothesis of highly-expressed testes genes generating a pattern of male over-expression in the top quartile. Furthermore, this pattern would be exacerbated on the Z chromosome due to the over-representation of male-biased genes on the Z, as observed here (see below) and also reported in *B. mori* (Arunkumar et al. 2009; Walters & Hardcastle 2011).

Finally, we note there is additional experimental evidence for a global sex chromosome dosage mechanism that down-regulates male Z expression, at least in *B. mori*. Kiuchi et al (2014) recently reported the characterization of a Z-linked *masculinizing* protein that plays a fundamental role in *B. mori* sex determination. RNAi disruption of this *masc* protein does not much alter autosomal expression, but causes chromosome-wide up-regulation of Z-linked expression. Thus *masc* apparently controls a switch that initiates a global epigenetic reduction in Z-linked male expression.

What does this current set of results from Lepidoptera mean for the *heterogametic dichotomy hypothesis*? These different lines of evidence pointing to a global sex chromosome dosage compensation mechanism in several lepidopteran species undermine the simple notion that female heterogametic taxa do not evolve such mechanisms. However, there is substantial room for nuance here. First, despite emerging support for Lepidoptera as an exception, there is still substantial evidence that global dosage compensation is much less common in ZW than XY taxa (Mank 2013). Thus, while seemingly not a universal rule, the *heterogametic dichotomy hypothesis* still reflects a compelling and unexplained trend in sex chromosome evolution.

Second, results presented here from *Heliconius* suggest there is a need to separately consider observed patterns, proximate mechanism, and evolutionary process. In most cases, reporting that a species shows a pattern of “incomplete” dosage compensation has been assumed to indicate the organism lacks a global mechanism to mitigate sex chromosome dosage differences between sexes. We now observe that *Heliconius* butterflies appear to break this assumption. *Heliconius* seems to share a mechanism with bombycoid moths that reduces male Z-linked expression in order to balance Z:A expression between sexes (Walters & Hardcastle 2011; Kiuchi et al. 2014; Smith et al. 2014). In bombycoids this balancing is “perfect”, resulting in “complete” sex chromosome dosage compensation with no male bias in Z-linked M:F expression ratios. Yet *Heliconius* shows a pattern of “incomplete” dosage compensation despite apparently deploying this mechanism of down-regulating the male Z. This result defies a simple characterization as being consistent, or not, with the *heterogametic dichotomy hypothesis*. On the one hand, opposing the hypothesis, *Heliconius* butterflies apparently have a global mechanism of dosage compensation. On the other hand, supporting the hypothesis, dosage effects persist and compensation is “incomplete”. A similar scenario was recently reported for the neo-X of *D. pseudoobscura*, where the well-characterized dosage-compensation complex appears in some tissues and developmental stages to be “incompletely” compensating male neo-X expression to an X:A ratio ∼0.85, much greater than 0.5 expected without compensation but still short of X:A ∼1 “complete” compensation observed in other *Drosophila* species and the ancestral portion of the *D. pseudoobscura* X (Nozawa et al. 2014). The functional and evolutionary significance of a lingering dosage effect in the presence of a global compensating mechanism is certainly a promising area for future research addressing the evolution of sex chromosome dosage compensation, especially the *heterogametic dichotomy hypothesis*. A reasonable argument can be made that reducing Z-linked expression in males to balance Z:A expression in females cannot legitimately be considered sex chromosome dosage compensation, *sensu stricto*. The canonical evolutionary model of dosage compensation invokes stabilizing selection to maintain ancestral expression levels of sex-linked genes in the heterogametic sex as gametologs erode from the W (or Y) (Ohno 1967; Charlesworth 1978; Mank 2009a). This model assumes that substantial reduction in the Z:A ratio is deleterious. Following this theory, sex chromosome dosage compensation should be defined as dosage *conservation* retaining the ancestral expression patterns where Z:A∼1. (This assumes all autosomes have roughly comparable average expression levels, a pattern that appears to be true in most species examined (Gupta et al. 2006; Deng et al. 2011; Hong Lin et al. 2011; Walters & Hardcastle 2011; Vicoso, Emerson, et al. 2013a; Vicoso, Kaiser, et al. 2013b).)

Thus, following this narrow and canonical view, sex chromosome dosage compensation should be limited only to situations where the heterogametic sex has Z:A ∼ 1 (and, comparably, X:A ∼1). Notably, this would include cases where sexual antagonism of sex-linked expression presumably remains unresolved and results in hyper-expression of sex-linked genes (i.e., X:A > 1) in the homogametic sex (Prince et al. 2010; Mank et al. 2011; Allen et al. 2013). However, a mechanism that globally reduces Z-linked expression in males to balance Z:A expression in females, as appears to exist in Lepidoptera, is not consistent with canonical evolutionary theory regarding dosage compensation and sex chromosomes. Specifically, if Z:A < 1 in heterogametic females is problematic, why would similarly reducing Z:A expression ratios in homogametic males resolve this problem? Doing so would compensate sex chromosome expression relative to autosomes between sexes, but presumably without dosage *conservation*. These results from Lepidoptera present a conundrum in light of current evolutionary theory (Walters & Hardcastle 2011).

Some resolution to this conundrum may come from better understanding the evolutionary history of the lepidopteran Z chromosome. Substantial ambiguity currently exists around the evolutionary relationship between the Z and W chromosomes. Basal Lepidoptera and Trichoptera (caddisflies, the sister group to Lepidoptera) are female-heterogametic but completely lack a W chromosome; females are ZO (Lukhtanov 2000; Sahara et al. 2011). The W chromosome now shared by most “advanced” (ditrysian) Lepidoptera arose long after the split between Lepidotera and Trichoptera. One hypothetical origin of the W invokes a fusion between an autosome and the basal Z chromosome, with the remaining free autosome following the canonical evolutionary degradation into the W allosome shared by ditrysian species (Traut & Marec 1996). In this case, standard expectations of dosage compensation apply and the current observations of Z:A < 1 in males remain theoretically inconsistent.

However, karyotypes of basal pre-ditrysian and ditrysian species do not obviously indicate a lost autosome, a fact consistent with an alternative scenario that posits the W originating as a supernumerary B chromosome that acquired a sex-determining locus and meiotic pseudo-bivalence with the Z (Lukhtanov 2000). In this latter scenario, origins of the Z chromosome are quite ancient, likely involve the transition to female-heterogamety, and existing theory offers few expectations about how sex chromosomes and dosage effects might evolve.

The available data indicate substantial variation among lepidopteran lineages in patterns of sex chromosome dosage compensation. Considered in light of the most recent lepidopteran phylogenies (Regier et al. 2013; Kawahara & Breinholt 2014), the patterns appear to reflect at least two evolutionary transitions in dosage compensation (Table 5). Two possible histories could explain these patterns. Either a similar dosage compensating mechanism evolved independently in butterflies and the ancestor of bombycoids, or an “incomplete” dosage compensating mechanism present in butterflies was refined to “completeness” in bombycoids but lost in pyralids.

**Table 5.**
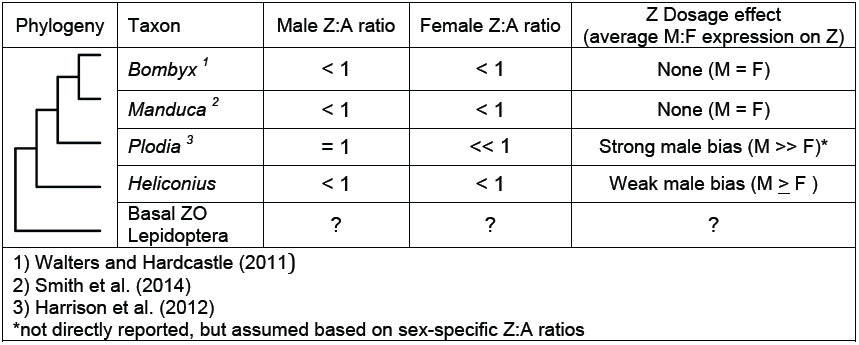
Summary of observed patterns of sex chromosome dosage compensation in the Lepidoptera.

| Phylogeny | Taxon | Male Z:A ratio | Female Z:A ratio | Z Dosage effect<br>(average M:F expression on Z) |
| --- | --- | --- | --- | --- |
|  | *Bombyx*<sup>1</sup> | < 1 | < 1 | None (M = F) |
|  | *Manduca*<sup>2</sup> | < 1 | < 1 | None (M = F) |
|  | *Plodia*<sup>3</sup> | = 1 | << 1 | Strong male bias (M >> F)* |
|  | *Heliconius* | < 1 | < 1 | Weak male bias (M ≥ F ) |
|  | Basal ZO<br>Lepidoptera | ? | ? | ? |
| 1) Walters and Hardcastle (2011)<br>2) Smith et al. (2014)<br>3) Harrison et al. (2012)<br>*not directly reported, but assumed based on sex-specific Z:A ratios |  |  |  |  |

Regardless of how dosage compensation has evolved among moths and butterflies, there are some intriguing similarities between the patterns seen among Lepidoptera and the emerging understanding of sex chromosome dosage compensation – or the lack thereof – in mammals. Although still controversial, there is a growing consensus the mammalian X evolved without dosage compensation and, in light of inferred ancestral expression levels on the X, the mammalian X:A ratio is ∼0.5 in both sexes (Julien et al. 2012; F Lin et al. 2012; Pessia et al. 2013). This view means that mammalian X-chromosome inactivation (XCI) effectively balances X-linked expression between sexes by halving the potential dose in females. A similar pattern was also recently reported in twisted-wing insects (Strepsiptera) (Mahajan & Bachtrog 2015). This scenario appears analogous to Lepidoptera, where Z-linked expression is similarly balanced between sexes by an apparent global reduction in expression in the homogametic sex. Investigation into the functional mechanisms of lepidopteran dosage compensation might help to resolve the origins of this mechanism. At this point it seems possible that Lepidoptera, Strepsiptera, and mammals have converged on similar mechanisms of mitigating sex chromosome dosage differences between the sexes.

### Chromosomal distribution of sex-biased genes

The general lack of sex-biased genes in purely somatic, non-reproductive tissue in *Heliconius* is unusual. There is good precedent from many other animals, particularly fruit flies and mice, that such tissues should contain at least dozens if not hundreds of genes with sex-biased expression (Yang et al. 2006; Catalán et al. 2012; Meisel et al. 2012). Even among other Lepidoptera such tissues yield numerous sex-biased genes (Walters & Hardcastle 2011; Smith et al. 2014). Thus it is tempting to explain away this result through a lack of statistical power or other methodological artifacts. Yet this seems unlikely because abdomens yielded thousands of differentially expressed loci. Heads and abdomens in this study were from the same individuals, with RNA extracted and analyzed simultaneously in parallel. Furthermore, the mouth, leg, and antennae data were generated independently of the head and abdomen samples but yielded a similar paucity of sex-biased genes. So it may simply be that *Heliconius* butterflies have unusually monomorphic patterns of gene expression between sexes for tissues unrelated to reproduction. Certainly phenotypic sexual dimorphism is quite limited in *Heliconius* compared to most moths and many butterflies (personal observations, JRW & CJD).

In contrast to non-reproductive tissues, the abdomens yielded a very large number of genes with significantly dimorphic expression. The distribution of such sex-biased genes is clearly different between the Z and autosomes, with male-biased genes more abundant and female-biased genes less abundant on the Z relative to autosomes, which have approximately equal amounts of male-and female-biased genes (Figure 4). A positive association between sex-linkage and expression biased towards the homogametic sex has been widely observed in several species of both male and female heterogametic taxa, including Lepidoptera (Reinke et al. 2000; Khil et al. 2004; Kaiser & Ellegren 2006; Arunkumar et al. 2009; Walters & Hardcastle 2011; Meisel et al. 2012). This pattern is generally predicted by models of sexually antagonistic evolution, specifically when sexually antagonistic mutations are fully or partially dominant (Rice 1984; Ellegren & Parsch 2007; Connallon & Clark 2010). Thus our results offer further support for the theory that the asymmetrical transmission of sex chromosomes favors sex-linked accumulation of mutations benefitting the homogametic sex.

A deficit of genes biased towards the heterogametic sex is also predicted by the same theory, but observations are much less consistent across taxa and seem to depend on a range biological idiosyncrasies including meiotic sex chromosome inactivation, tissue specificity of expression, and mechanisms of dosage compensation (Connallon & Clark 2010; Meisel et al. 2012; Parsch & Ellegren 2013). Nonetheless, there is at least one promising coincidence of theory and data that has been recently noted: species with equal rates of recombination between sexes tend to show an excess of sex-linked genes with heterogametic bias (e.g. birds, mammals), while in *Drosophila*, where males do not recombine, a paucity of male-biased genes on the X is often reported (Connallon & Clark 2010). Since lepidopteran females lack recombination, the paucity of female-biased genes on the Z in *Heliconius* is consistent with this pattern. However, our data represent only an initial and broadly scoped assessment of this phenomenon.

## Conclusion

Our results are consistent with an increasing body of evidence that many species of moths and butterflies possess a sex chromosome dosage compensating mechanism that operates by reducing Z chromosome expression in males. However, this mechanism appears to be imperfect in *Heliconius*, resulting in a moderate Z chromosome dosage effect. These results counter the emerging view that female-heterogametic ZW taxa have incomplete dosage compensation because they lack a chromosome-wide epigenetic mechanism mediating sex chromosome dosage compensation. In the case of *Heliconius*, sex chromosome dosage effects apparently persist despite such a mechanism. This result adds additional complexity to patterns of dosage compensation observed in Lepidoptera. Finally, patterns of sex-biased expression in our data highlight the impact of sexually antagonistic selection in shaping genome evolution.

## Acknowledgements

This research was supported in part by a NSF post-doctoral fellowship to JRW (DBI-0905698). RNA sequencing was funded by the “Capacity and Capability Challenge Programme” from The Genome Analysis Centre, Norwich, UK. Luiqi (Aloy) Gu provided valuable comments on the manuscript.

## Appendix Analysis of Z-linked expression quartiles between male and female

Harrison et al. (2011) analyzed patterns of sex-chromosome dosage compensation in the Indian meal moth, *Plodia interpunctella.* One aspect of their analysis contrasted male and female expression of Z-linked loci split by quartiles of expression magnitude. This analysis is intended to illustrate that dosage effects increase with gene expression level. Importantly, Harrison et al. defined quartiles of expression based solely on male expression; female expression was ignored when binning loci into subsets to assess dosage effects. Here we demonstrate that, in the context of RNA-seq data, their approach potentially leads to an artifactual bias that would result in expression appearing greater in males than females for highly expressed loci, as reported by Harrison et al. (2011).

We judge it unlikely that this bias can completely explain the patterns reported for *P. interpunctella.* Therefore we have not re-analyzed the data of Harrison et al. and we do not question their qualitative conclusions regarding the relationship between relative dosage effects and the magnitude of gene expression. Nonetheless, for the sake of future analyses of sex chromosome dosage compensation, we feel it worthwhile to describe what we believe to be a better method of anlaysis. We suggest that quartile binning based on the *maximum of male or female expression at each locus* provides a simple adjustment to the analysis that removes bias imposed by binning on data from only one sex.

This analysis tests whether dosage effects (i.e. greater Z-linked expression in males) increases with magnitude of gene expression. It is based on the idea that, for any given locus, the maximum level of gene expression is limited by the rate of translation because the translational machinery becomes saturated. In the absence of sex chromosome dosage compensation, highly expressed genes at or near the point of maximum translation will exhibit a strong dosage effect; the single female gene copy cannot reach the transcription rate of two copies in male. In contrast, genes expressed at low to intermediate levels will exhibit little or no dosage effect because sufficient transcripts can be generated from a single female copy to balance transcript levels in males. The resulting pattern is a discrepancy between male and female Z-linked expression that increases with the magnitude of gene expression.

In principle, it should be possible capture this effect by comparing the distributions of male and female expression of Z-linked genes that are expressed at relatively high or low absolute expression levels. Following the approach of Harrison et al., binning loci by quartiles of expression should reveal increasingly greater relative expression in males with each additional quartile if translational saturation effects are occurring in the absence of dosage compensation.

However, *it is in the grouping of loci into bins of relative expression levels where a bias may arise.* Binning based on expression in only one sex can cause expression in that sex to appear relatively reduced among weakly expressed genes, but relatively high for strongly expressed genes, even if the distribution of gene expression is identical for both sexes. Thus grouping loci by expression quartiles using values in only one sex can cause biases in which sex appears to have relatively greater expression in each quartile. This phenomenon is best demonstrated with a simple simulation in which the average expression of males and females is identical.

Described briefly, the simulation begins by generating “true” expression values for 10,000 loci as random draws from a log-normal distribution. Variation in expression levels for male and female “observations” are created by independently adding values drawn from a normal distribution with a mean of 0 to each “true” expression value. Additionally, 1000 loci are further modified to be differentially expressed between males and females, 500 in each direction. For the sake of brevity and clarity, we refer the reader seeking further details to the attached R script encoding the simulation. The “MA-plot” of simulated gene expression data (Figure A1) should suffice to convince most readers that the simulated data reasonably represents gene expression data.

**Fig. A1.**
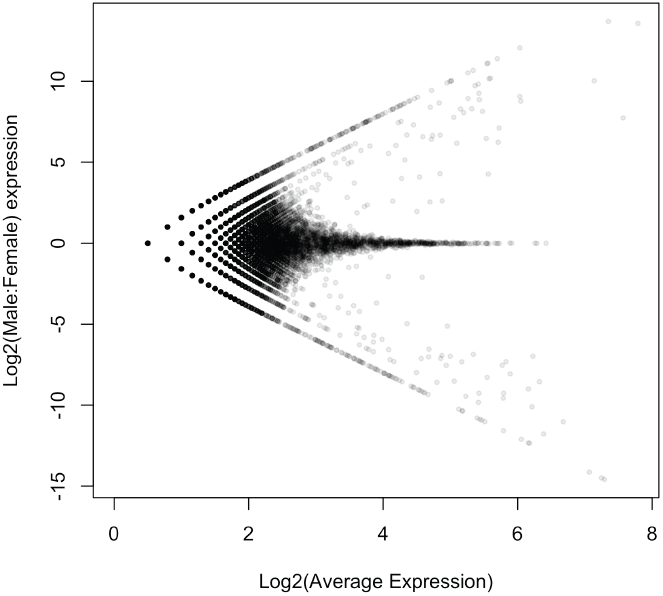
MA-plot of simulated male and female expression data.

Using these simulated values, we can split the dataset into quartiles of expression and compare the distributions between sexes (Figure A2). Intuitively, since all differences between male and female expression are random and unbiased, we would expect no systematic differences between sexes at any expression level. However, this outcome is only observed if quartiles are based *on the maximum of male or female expression* for each locus. Ranking using only one sex clearly results in a strongly biased outcome that depends on which sex is used for ranking (Figure A2).

**Fig. A2.**
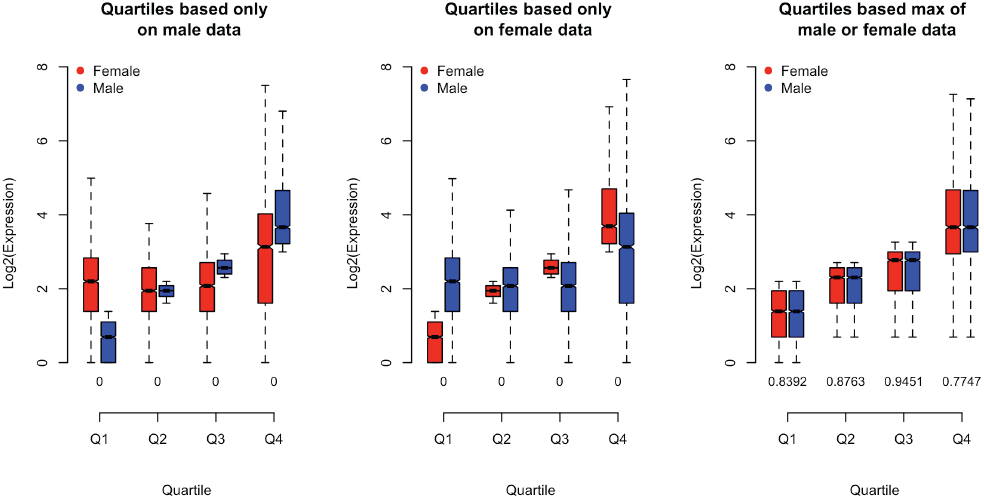
Quartile comparisons of simulated gene expression distributions of simulated male and female expression levels. Comparison of male (blue) and female (red) expression levels for loci, divided into quartiles based on male expression (left panel), female expression (center panel), and the maximum male or female expression (right panel). Boxes indicate the interquartile range (IQR) around the median (black bar) and whiskers extend to 1.5 times the IQR. P-values of a Mann-Whitney U test of inequality between males and females are printed for each quartile along the x-axis, with “0” indicating P < 0.0001.

We emphasize that this simulation has been created primarily to illustrate our point about biases that potentially arise from quartile analyses based on ranking on one sex alone. We do not claim that the simulation is highly realistic and we have not extensively explored the effect of changing simulation parameters. Our narrow aim is to communicate an intuitive and straightforward demonstration of why we have deviated from the precedent set by Harrison et al. (2012) for this type of analysis.

---

**Figure S1.**
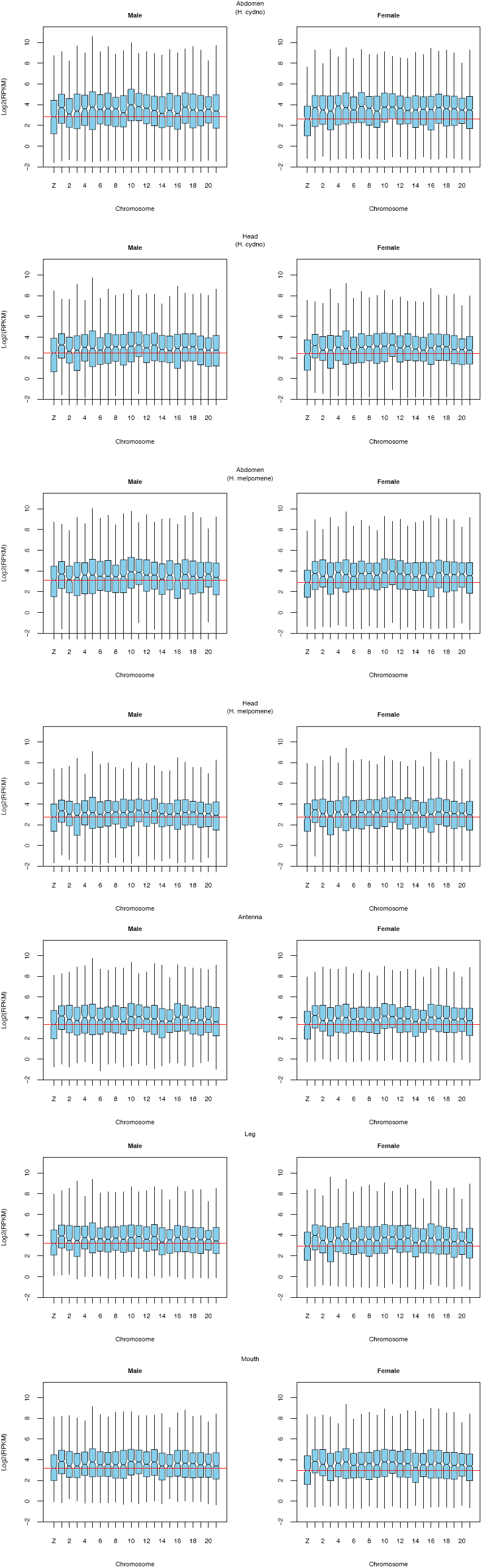
Distributions of log2-transformed gene expression levels (FPKM) for each ome in male and female Heliconius butterflies. The right-hand most box reflects t mapped to chromosome. The horizontal redline indicates the median log2(FPKM) of the Z chromosome. Boxes indicate the interquartile range (IQR) around the black bar) and whiskers extend to 1.5 times the IQR; outlier points beyond 1.5 IQR lotted. Data were filtered with a null expression likelihood of 0.5.

**Fig S2.**
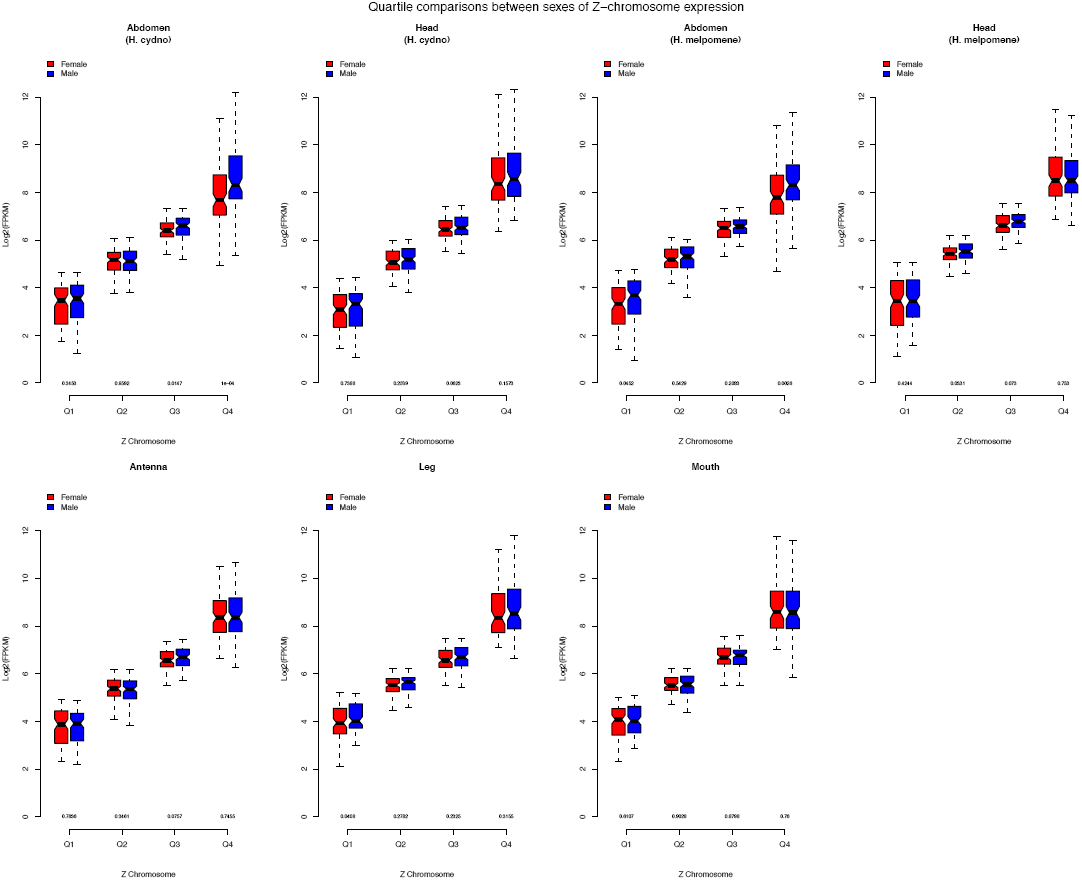
Comparison of male (blue) and female (red) expression levels for Z-linked loci, divided into quartiles based on the maximum of male or female expression in various tissues from Heliconius butterflies. Boxes indicate the interquartile range (IQR) around the median (black bar) and whiskers extend to 1.5 times the IQR. P-values of a Mann-Whitney U test of inequality between males and females are printed for each quartile along the x-axis.

**Figure S3.**
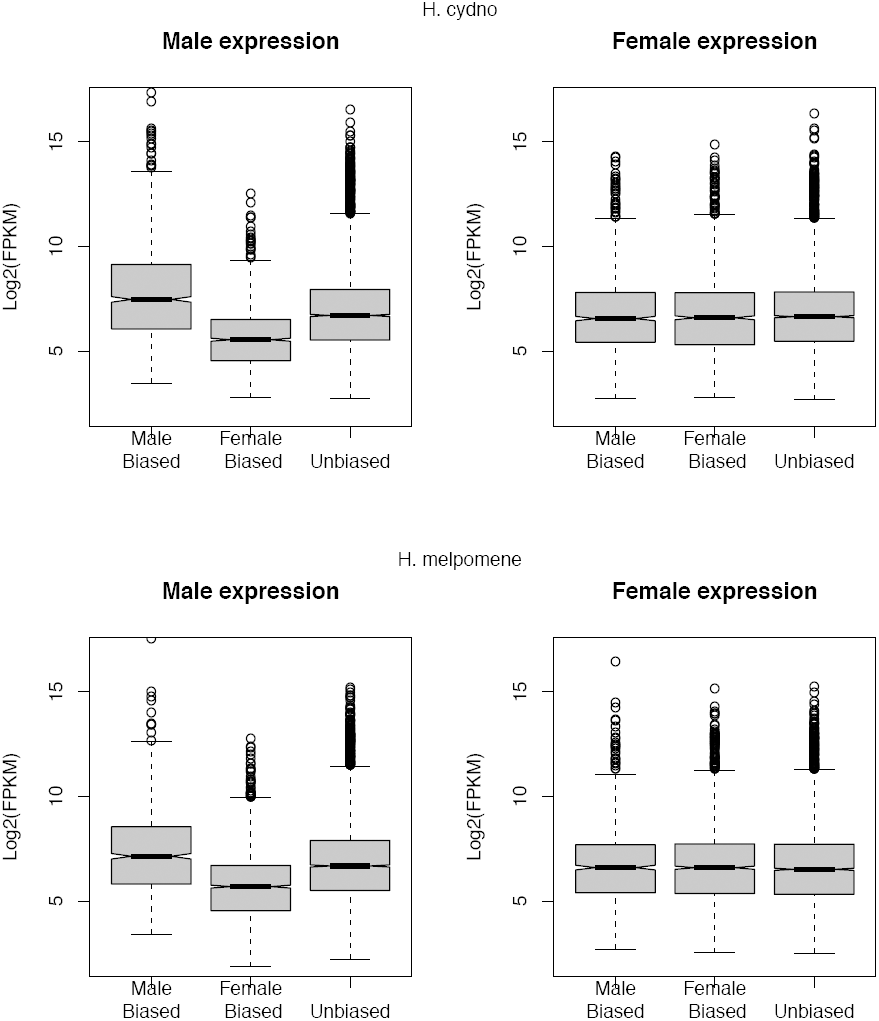
Distributions of log2-transformed gene expression levels (FPKM) for expressed autosomal loci, partitioned by category of sex-biased expression, in abdomens from two species of Heliconius butterflies. In both species males, but not females, showed significantly different levels of expression across categories (Kruskal-Wallis test, p ≪ 0.001). A gene was considered sex-biased if significantly differentially expressed between males and females with FDR < 0.05 and a minimum fold-change of 1.5. Boxes indicate the interquartile range (IQR) around the median (black bar) and whiskers extend to 1.5 times the IQR. Data were filtered with a null expression likelihood of 0.5.

